# From default to creativity: prefrontal and cerebellar contributions of the default mode network to goal-directed remote thinking

**DOI:** 10.64898/2026.03.14.711790

**Authors:** Victor Altmayer, Sarah Moreno-Rodriguez, Marcela Ovando-Tellez, Benoît Béranger, Alizée Lopez-Persem, Emmanuelle Volle

## Abstract

Creativity is a hallmark of human cognition, characterized by the ability to connect seemingly distant concepts or ideas. Existing theories suggest that remote thinking can be achieved either spontaneously (constrained by the structure of semantic memory) or in a goal-directed manner (constrained by a creative goal). The present study investigates the neural correlates of goal-directed remote thinking, defined as the intentional production of semantically distant associations.

Using a simple word-to-word association task comprising both a spontaneous condition and a goal-directed creative condition, we investigated *Goal-directed Remoteness* as the extra semantic distance traveled away from spontaneous responses when instructed to think creatively. Task-based functional MRI in 38 healthy young adults identified brain regions whose activation scaled with *Goal-directed Remoteness*. The results revealed that activity in the rostromedial and dorsomedial prefrontal cortex and the right cerebellar Crus I & II was positively modulated by *Goal-directed Remoteness.* Control analyses confirmed the robustness of these findings independently of the cue-words’ semantic or linguistic properties. Follow-up seed-based resting-state functional connectivity analyses characterized the intrinsic connectivity profiles of the revealed regions. They showed that rostromedial and dorsomedial prefrontal and cerebellar Crus I & II regions formed a functionally interconnected network primarily overlapping with the default-mode network (DMN).

Our findings challenge the traditional view of the DMN as supporting only passive or spontaneous cognition. Instead, they reveal a prefronto-cerebellar DMN subnetwork supporting goal-directed remote thinking, a key component of creative cognition. Within this network, the rostromedial and dorsomedial prefrontal cortex and cerebellar Crus I & II play an active role in the intentional generation of connections between distant concepts.

## Introduction

What is the first word that comes to your mind when you think of the word “mother”? Now, think of another word associated with “mother” in a more creative way. Executing this type of task relies on remote thinking, a core component of creativity(Bendetowicz et al. 2018; Green et al. 2015; Prabhakaran et al. 2014; Mednick 1962), allowing people to link mutually distant semantic concepts. For instance, the uncommonness or remoteness of responses in word-to-word generation tasks has been shown to correlate with creative performance.(Altmayer et al. 2024; Bendetowicz et al. 2018; Green et al. 2015; Prabhakaran et al. 2014) Remote thinking is central to creative thinking, not only in verbal but also domain-general creativity.(Mednick 1962; Beaty et Kenett 2023) Hence, clarifying the neurocognitive processes of remote thinking may help to understand some of the essential mechanisms of idea generation during creative thinking.(Beaty, Kenett, et al. 2018; Beaty et al. 2021; 2016; 2015)

The generation of creative semantic associations was historically thought to rely on spontaneous associative thinking processes that connect seemingly unrelated concepts.(Mednick 1962; Volle 2018; Beaty et Kenett 2023; Benedek et al. 2023) Associative thinking is classically considered a spontaneous, uncontrolled process shaped by individuals’ semantic memory structure, whose variations can account for individual differences in creativity.(Beaty et Kenett 2023; Luchini et al. 2023; Ovando-Tellez et al. 2022; He et al. 2020; Benedek et Jauk 2018; Kenett 2018; Kenett et al. 2014) Traditionally linked to spontaneous, unconstrained cognitive processes(Bar et al. 2007; Raichle 2015), the default mode network (DMN) has been proposed to support spontaneous aspects of creative cognition and remote thinking.(Volle 2018; Bendetowicz et al. 2018; Marron et al. 2018; Beaty et al. 2016)

However, recent findings suggest that remote associative thinking is not always spontaneous and can also be constrained by a creative goal.(Benedek et Fink 2019; Green et al. 2024) Hence, goal-directed aspects of remote thinking may complement a strictly spontaneous propagation within a semantic structure during creative thinking.(Altmayer et al. 2024; Benedek et al. 2023; Beaty et Kenett 2023) Indeed, because spontaneous associations are often unoriginal, creativity often requires the intentional search for remote semantic associates. This deliberate activation of mutually remote semantic concepts, or “goal-directed remote thinking”, is thought to play a key role in both divergent (requiring the generation of many original ideas) and convergent (requiring the constrained search for a unique but remote solution to a problem) aspects of creativity by enabling individuals to move beyond conventional associations and generate original ideas.(Prabhakaran et al. 2014; Green et al. 2015; Bendetowicz et al. 2018; Altmayer et al. 2024) Furthermore, if we adopt the position of Green et al.(Green et al. 2024), who recently defined creativity as an “internally-oriented attentional process with a generative goal”, then goal-directed remote thinking may represent the core component of creative cognition.

The classical dual process framework assumes that during creative thinking, executive control processes supported by the executive control network (ECN)(Seeley et al. 2007; Power et Petersen 2013) interact with spontaneous associative processes to combine, elaborate, manipulate, evaluate, select, or reject spontaneously generated semantic associations depending on the goal(Benedek et Jauk 2018; Volle 2018; Beaty et al. 2016). Such executive control processes include inhibition, updating, and flexibility, but their exact role in goal-directed remote thinking remains unclear. In a recent study, Altmayer et al.(Altmayer et al. 2024) isolated goal-directed remote associative processes from cognitive inhibition, both behaviorally and in the brain. Their results suggested that goal-directed remote thinking and executive control processes have distinct brain substrates and may have distinct roles in thinking creatively. Beyond the creativity field, the generation of remote semantic associations has been studied under the concept of controlled retrieval and has been found to involve regions of the semantic control network and DMN(Jefferies 2013; Lambon Ralph et al. 2017; Krieger-Redwood et al. 2023), in particular the left inferior frontal gyrus and the dorsomedial prefrontal cortex (PFC).(Badre et al. 2005; Hirshorn et Thompson-Schill 2006; Krieger-Redwood et al. 2023) Finally, semantic goal representation was shown to involve ECN but also DMN regions.(Wang et al. 2021) Hence, separate lines of research suggest distinct brain candidates possibly supporting goal-directed remote thinking, including ECN, semantic control network, and DMN regions.

In this study, we aimed to identify the brain regions involved in goal-directed remote thinking, measured as the ability to think away from obvious, spontaneous, free word associations. Goal-directed remote thinking was operationalized in the Free Generation of Associates Task (FGAT)(Bendetowicz et al. 2018; Lopez-Persem, Moreno-Rodriguez, et al. 2024; Moreno-Rodriguez et al. 2025; Battistello et al. 2025; Altmayer, Ovando-Tellez, Bieth, Batrancourt, Rametti-Lacroux, Bernardaud, et al. 2025; Altmayer, Ovando-Tellez, Bieth, Batrancourt, Rametti-Lacroux, Moreno-Rodriguez, et al. 2025), a simple word-association task, in which participants are asked to provide both spontaneous and intentionally creative word-to-word associations. While asking participants to think creatively may appear simplistic, it has been shown to positively influence creative thinking and to relate to other measures of creativity.(Prabhakaran et al. 2014; Bendetowicz et al. 2018; Wei et al. 2024) Young healthy participants performed the FGAT during fMRI. Parametric modulation analyses allowed us to identify brain regions whose activation varied with the semantic remoteness of goal-directed remote associations compared to spontaneous associations. To further characterize each of the identified brain regions, we then applied seed-to-voxel resting state functional connectivity (RSFC) analyses to reveal each region’s intrinsic connectivity network.

## Materials and methods

### Participants

Forty participants were recruited and tested at the CENIR core facility of the Paris Brain Institute (ICM). All participants were French native speakers, right-handed, with correct or corrected vision, and free from neurological or psychiatric disorders. Two participants were excluded from analyses because of MRI scanning interruptions: one because of a claustrophobic episode and the other because of a technical issue, resulting in a final sample of thirty-eight participants (19 females, age = 26.5 ±4.0 (M±SEM), years of education = 16.5 ±1.9 (M±SEM)). All were paid 110€ for a four-hour testing session including MRI scanning and cognitive and creativity tests inside and outside the scanner. The current study builds upon a recently published dataset whose analyses focused on valuation processes involved in creative thinking.(Moreno-Rodriguez et al. 2025) Instead, the current work focuses on the brain correlates of remote thinking during the FGAT, which were not examined before. An approved ethics committee authorized the study (CPP Ouest II – Angers), all participants provided written informed consent, and confidentiality was preserved in accordance with French regulations.

### Free generation of associates task (FGAT)

To explore remote thinking, we used the FGAT, a simple word-association task previously validated in patients(Bendetowicz et al. 2018; Altmayer et al. 2024; Altmayer, Ovando-Tellez, Bieth, Batrancourt, Rametti-Lacroux, Moreno-Rodriguez, et al. 2025) and healthy subjects.(Lopez-Persem, Moreno-Rodriguez, et al. 2024; Moreno-Rodriguez et al. 2025; Battistello et al. 2025)

#### Experimental design

The FGAT aims to distinguish goal-directed from spontaneous semantic associations and is thus divided into two successive conditions, each comprising 5 training trials and 62 test trials.

The ‘FGAT-First’ condition explores spontaneous associative thinking.(Mednick 1962) In this condition, participants were asked to provide the first word coming to mind in response to a cue-word (e.g., “father” in response to the cue “mother”).

In the ‘FGAT-Distant’ condition, participants were asked to intentionally generate a word associated with the cue-word in an original, creative way, while still being understandable to someone else (e.g., “nature” in response to the cue “mother”). This condition explores intentional remote thinking, thought to involve both spontaneous and goal-directed associative processes.

Participants always performed the ‘FGAT-First’ before the ‘FGAT-Distant’, with the same 62 cue-words for both conditions, successively presented in a randomized order. Before the task began, the instructions were displayed on a screen, and the examiner clarified that responses should be a single word, not a phrase, a compound word, or a proper noun. Each trial displayed a fixation cross for 1.6 to 3.2 seconds, jittered with a uniform distribution. Then, a cue-word was displayed on the screen, and participants were asked to press a response button with their right index finger when they had an answer in mind, before saying their response aloud with a microphone. The experimenter typed the response and displayed it on the participant’s screen. Participants could then repeat their response in case the experimenter had misheard (by pressing the response button again with their right index finger) or to validate the response (by pressing a validation button with their right ring finger). The cue-words were displayed until participants produced an oral response, within a time limit of 10 seconds for the ‘FGAT-First’ condition, and 20 seconds for the ‘FGAT-Distant’ condition.

We used MATLAB (Version 9.9.0.1495850, R2020b; The MathWorks Inc., Natick, MA, USA) to program the task and to extract variables of interest.

#### FGAT material

To build the verbal material for the FGAT, we used French nouns with a mean lexical frequency of 102 occurrences per million according to mean written frequencies (Supplementary Table 1; text- and web-based word frequencies computed with Lexique 3.8(New et al. 2004); www.lexique.org). As cue-words may vary in their strength of association with other words, which may impact participants’ responses, we quantified the semantic richness(Beaty et al. 2022) and associative steepness(Mednick et al. 1964) of the FGAT cue-words, and used them as covariates in control analyses.

We quantified the semantic richness of each cue-word as the ratio between the number of different semantic associates for this cue-word and the total number of responses given for this cue-word, based on the French free-association norms.(Debrenne 2011) This ratio indicates the relative number of semantic associates linked to a given cue-word that compete for selection, and thus may indicate the degree of interference the participant must overcome when searching for and selecting a response. The number of such competitors may negatively impact originality.(Beaty et al. 2022) The semantic richness of the 62 cue-words ranged from 8.48 to 34.32, with a mean value of 19.21 (Supplementary Table 1).

We also quantified cue-words’ steepness. A steep cue-word (e.g., “cat”) is one for which there is a much stronger semantic association to one associate (“dog”) than to all other associates. Thus, steep cue-words induce strong associative constraints during FGAT, and require inhibition of this strong first associate to find more original associations in the distant condition. This measure is also relevant as, according to Mednick’s theory, more creative people have flatter hierarchies in their associations stored in memory.(Mednick 1962) Steepness is computed as the frequency ratio between the most frequent and the second most frequent associates of a given cue-word. The 62 cue-words used in the FGAT were selected to ensure equal proportions of steep (steepness > 4) and flat (steepness ≤ 3) cue-words.(Bendetowicz et al. 2018) Associative steepness ranged from 1.03 to 24.08, with a mean value of 4.83 (Supplementary Table 1).

#### FGAT scoring

All responses were first manually corrected for spelling errors, typos, and special characters, and homogenized responses’ declension forms. Only valid responses (i.e., simple words excluding proper nouns) were considered for further analysis.

For each trial in each condition, we computed the semantic similarity between the response and the cue-word using a pre-trained word2vec model for French built from a 1.6 billion-words corpus constructed from websites with .fr domains (https://fauconnier.github.io,(Fauconnier 2015)). Word2vec models are word-embedding algorithms based on neural networks, computing vector representations of words in a text so that words that share similar contexts are represented by close numerical vectors.(Mikolov et al. 2013) They can be used to compute semantic similarity between cue-words and responses, from −□1 (low similarity) to 1 (high similarity). For each cue-word, we then calculated *Goal-directed Remoteness* as the difference in semantic similarity between ‘FGAT-First’ and ‘FGAT-Distant’ responses, allowing us to explore how much more remote from the cue-word the responses in the ‘FGAT-Distant’ condition were compared to those in the ‘FGAT-First’ condition. This score thus reflects the extra semantic distance individuals travel when their goal is to be creative.

### Task-based functional MRI

#### MRI Acquisition

MRI acquisitions were performed on a 3T Siemens Magnetom Prisma Fit MRI scanner with a 64-channel total imaging matrix system. We acquired two fMRI runs for the two task conditions (FGAT-First and FGAT-Distant) and one resting-state fMRI (rs-fMRI) run of 10 minutes (360 volumes). For each run we used multi-echo echo-planar imaging (EPI) sequences (repetition time (TR) = 1660 ms; echo times (TE) for echo 1 = 14.2 ms, echo 2 = 35.39 ms, and echo 3 = 56.58 ms; flip angle = 74°; 60 slices, slice thickness = 2.50 mm, iso-voxel; Ipat acceleration factor = 2; multiband = 3; and interleaved slice ordering). Because of the self-paced experimental design, the number of volumes per task fMRI run varied between FGAT conditions and between participants (mean, [min; max]: FGAT-First= 379, [317;454], FGAT-Distant: 612, [359;856]). We did not record any dummy scans and, therefore, did not discard any volume. After the FGAT participants also performed another task in the scanner that was the focus of a previous study(Moreno-Rodriguez et al. 2025), but out of the scope of the current one. In the current study, we focus on the FGAT-Distant condition. We also acquired high-resolution T1-weighted structural images (TR = 2300 ms; TE = 2.76 ms; flip angle 9°; 192 sagittal slices with a 1-mm thickness, iso-voxel; Ipat acceleration factor = 2).

#### Preprocessing

We used the afni_proc.py pipeline from the Analysis of Functional Neuroimages software (AFNI; https://afni.nimh.nih.gov) to preprocess all fMRI runs. Preprocessing steps included slice-timing correction and realignment to the first volume (computed on the first echo). Preprocessed data were then combined using TE-dependent analysis of multi-echo fMRI data (TEDANA; https://tedana.readthedocs.io/, version 0.0.9a1(DuPre et al. 2021)). We then used Statistical Parametric Mapping (SPM) 12 package running in MATLAB (MATLAB R2017b, The MathWorks Inc., USA) to co-register the resulting data onto the T1-weighted structural image, before normalizing the data to the Montreal Neurological Institute (MNI) template brain using the transformation matrix computed from the normalization of the T1-weighted structural image with the computational anatomy toolbox (CAT 12; http://dbm.neuro.uni-jena.de/cat/, implemented in SPM(Gaser et al. 2024)).

#### Task fMRI analysis

To analyze the resulting normalized task-related fMRI data, we used a SPM general linear model (GLM) with 24 motion parameters and 42 physiological nuisance regressors. These were computed using the Matlab PhysIO Toolbox (version 5.1.2, TAPAS software collection: https://www.translationalneuromodeling.org/tapas(Frässle et al. 2021). These nuisance regressors involved (i) physiological (respiration, cardiac pulse) noise recordings used to generate RETROICOR(Glover et al. 2000) regressors, (ii) White Matter (WM) and CerebroSpinal Fluid (CSF) masks from the anatomical segmentation used to extract components from compartments of non-interest using a Principal Component Analysis (PCA), and (iii) motion parameters, composed of standard motion parameters, their first temporal derivatives, and quadratic terms of each parameter and each temporal derivative. Additionally, time series were smoothed with an 8 mm full-width at half maximum Gaussian kernel, and frames exceeding a threshold of 0.5 mm Framewise Displacement were removed.

We then applied the GLM on the pre-processed FGAT-Distant time series to explore the neural encoding of *Goal-directed Remoteness.* The model included a boxcar function capturing the signal during the trial, i.e., between the cue display (onset) and the response button press (one event per trial), with *Goal-directed Remoteness* as a parametric modulator. To control for multiple comparisons, results were corrected using a FWE rate at *P* < .05 at the cluster level.

Control analyses employed similar GLMs after orthogonalizing *Goal-directed Remoteness* by cue-words’ richness, steepness, or trials’ response time, to ensure the stability of the results across variations in cue-words’ semantic properties and response time.

#### Seed-based resting-state fMRI analysis

We then used a seed-based approach to explore the intrinsic functional connectivity of significant clusters identified in the task-based fMRI analysis. Using Nilearn 0.9.2 (nilearn.github.io) running on Python 3.10.4 (www.python.org), BOLD time series underwent z-scoring, smoothing with a 2 mm full-width at half maximum Gaussian kernel, detrending, band-pass filtering within the 0.01–0.1 Hz range, and removal of frames exceeding 0.5 mm Framewise Displacement. For each significant cluster from the task fMRI analysis, taken as region-of-interest (ROI), and each participant, we computed ROI-to-voxel rs-fMRI connectivity maps correlating the mean fMRI time series within each ROI with the time series of each brain voxel. Fisher’s z-transform was applied to improve the normality of ROI-to-voxel connectivity maps. To explore the group-level connectivity profile of each ROI, we then conducted one-sample, one-tailed t-tests in SPM of z-transformed ROI-to-voxel correlation maps across participants. To control for multiple comparisons, results were corrected using a FWE threshold at *P* < .05 at the cluster level with a spatial cluster extent threshold of 10 contiguous voxels. To interpret the functional connectivity patterns of each ROI, we examined the overlap of each ROI’s connectivity with Yeo and colleagues’ referential intrinsic functional connectivity parcellations in 7 large-scale brain networks.(Yeo et al. 2011; Buckner et al. 2011; Choi et al. 2012)

### Other statistical analyses

The difference in semantic similarity between FGAT-First and FGAT-Distant conditions was tested using a Wilcoxon rank paired t-test.

We used separate linear mixed-effects models with random effects on both individuals and cue words to explore whether cue-words’ semantic richness, steepness, or trial response time predicted *Goal-directed Remoteness*.

## Results

### Behavioral results

As expected from the task design, the semantic similarity with the cue-word was higher for FGAT-First (0.36 ± 0.25 (M±SEM)) responses than for FGAT-Distant (0.18 ± 0.18 (M±SEM)) responses (*W* = 2.18 x10^6^; *z* = 25.88; *P* < 0.001) (Supplementary Figure 1).

Using linear mixed-effects models (Supplementary Table 2), we found a significant positive effect of cue words’ steepness on *Goal-directed Remoteness* (β =1.99 x10^-2^, *SE* = 6.24 x10^-3^, *t*(52.68) = 2.97, *P* = .004), and a significant negative effect of cue words’ semantic richness on *Goal-directed Remoteness* (β = -1.36 x10^-2^, *SE* = 2.46 x10^-3^, *t*(28.00) = -5.54, *P* < .001), while no significant effects was found for FGAT-Distant trials’ response time (β = 8.88 x10^-7^, *SE* = 2.05 x10^-6^, *t*(37.94) = 0.43, *P* = 0.668). Thus, cue words with higher steepness (i.e., a stronger dominant semantic associate) were associated with greater *Goal-directed Remoteness*, meaning that intentionally remote associations more semantically distant from the cue word than spontaneous associations. On the other hand, cue words with higher semantic richness were associated with a reduced *Goal-directed Remoteness*.

### *Goal-directed Remoteness*-related brain activation

To explore which neural regions encode the ability to intentionally provide semantic associations that are more creative than spontaneous ones, we first examined the relationship between *Goal-directed Remoteness* and neural activity during the FGAT-Distant task. A whole-brain parametric modulation approach revealed that activity in the bilateral dorsomedial and rostromedial PFC, and Crus I & II of the right cerebellum (Table 1, Figure 1) was scaled by *Goal-directed Remoteness*.

**Figure 1.**
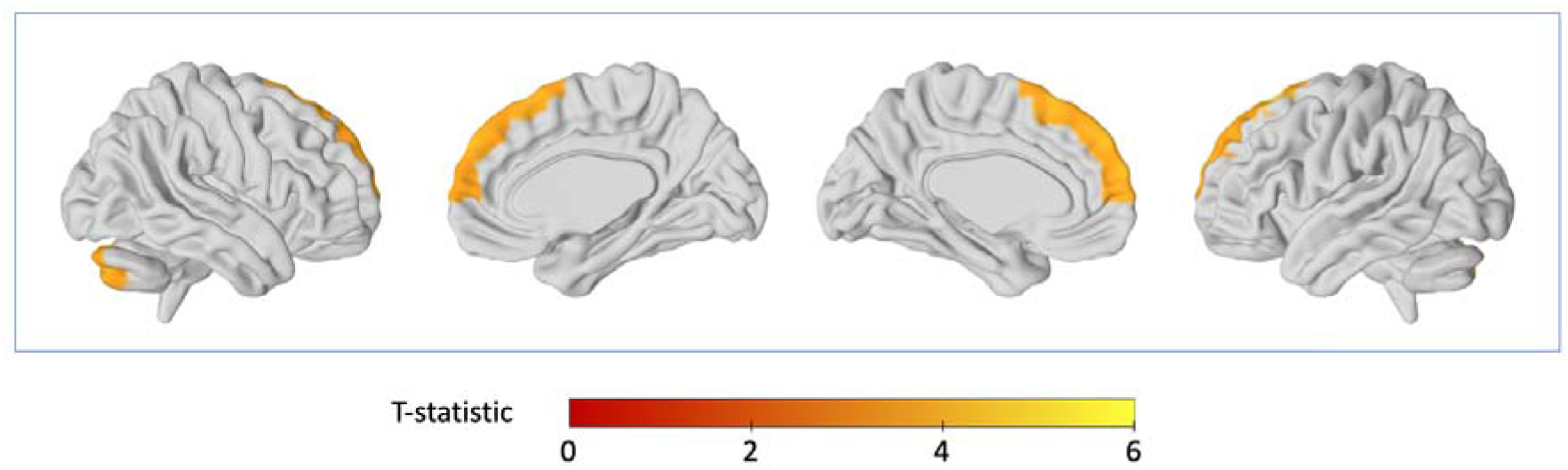
*Goal-directed Remoteness-*related brain activation. Whole-brain parametric modulation revealing significant regions modulated by *Goal-directed Remoteness* during FGAT distant. Significant clusters are shown in color. Statistical analyses were thresholded for significance at *P_FWE_* < 0.05 at the cluster level with a spatial cluster extent threshold of 10 contiguous voxels.

**Table 1.** Significant peak coordinates of the whole-brain parametric modulation by Goal-directed Remoteness. Significant clusters from the whole-brain fMRI analysis of the FGAT-Distant task with Goal-directed Remoteness as parametric modulator are reported, including MNI coordinates, T-score and Puncorrected value of significant clusters’ maxima, cluster size, cluster PFWE value, and related Brodmann areas. Statistical analyses were thresholded for significance at Puncorrected < 0.001 at the voxel level and PFWE < 0.05 at the cluster level.

| Brain region | BA | MNI peak coordinates |  |  | Peak level |  | Cluster level |  |
| --- | --- | --- | --- | --- | --- | --- | --- | --- |
| | | x | y | z | T | $P_{uncorrected}$ | voxels | $P_{FWE}$ |
| Bilateral dorsomedial PFC | 6 | **-8** | **25** | **60** | 5.57 | 0.000 | **227** | **0.002** |
|  |  | 6 | 25 | 63 | 4.59 | 0.000 |  |  |
|  |  | 2 | 12 | 68 | 3.73 | 0.000 |  |  |
| Bilateral rostromedial PFC | 10, 9 | **-8** | **62** | **23** | 5.18 | 0.000 | **346** | **0.000** |
|  |  | -6 | 55 | 38 | 5.06 | 0.000 |  |  |
|  |  | 6 | 58 | 38 | 4.87 | 0.000 |  |  |
| Right cerebellum, Crus I & II | NA | **29** | **-78** | **-37** | 4.94 | 0.000 | **133** | **0.029** |

BA: Brodmann Areas; FWE: Family Wise Error; MNI: Montreal Neurological Institute; NA: Not Applicable; PFC: Prefrontal Cortex

Control analyses after orthogonalizing *Goal-directed Remoteness* on cue-words’ semantic richness, steepness, or trials’ response time revealed similar results (Supplementary Tables 3 to 5, Supplementary Figures 2 to 4), identifying the same three clusters as in the original analysis, except for the right cerebellum, which was not significantly modulated by *Goal-directed Remoteness* after orthogonalization on cue-words’ steepness.

### Intrinsic connectivity of *Goal-directed Remoteness*-related brain regions

Using a seed-based approach, we then investigated the intrinsic connectivity profile of each of the three brain regions identified as activated during *Goal-directed Remoteness*. Each region was shown to be connected to the two other regions, forming a functional network largely overlapping with the DMN (Figure 2, Table 2). In addition to reciprocal interconnections between these three regions, the rostromedial PFC cluster was connected to other key nodes of the DMN, namely the bilateral precunei, posterior cingulate, inferior parietal, and anterolateral temporal cortices. This rostromedial PFC’s intrinsic connectivity network overlapped at 80% with the DMN. Cerebellar Crus I & II were also found connected to the bilateral precunei and posterior cingulate cortices, with a 75% overlap of their intrinsic connectivity network with the DMN. Cerebellar ROIs were also connected with local cerebellar regions overlapping with the ECN (accounting for 23% of their intrinsic connectivity). Finally, the dorsomedial PFC also connected to the right paracentral cortex, and showed a 70% overlap of its intrinsic connectivity network with the DMN.

**Figure 2.**
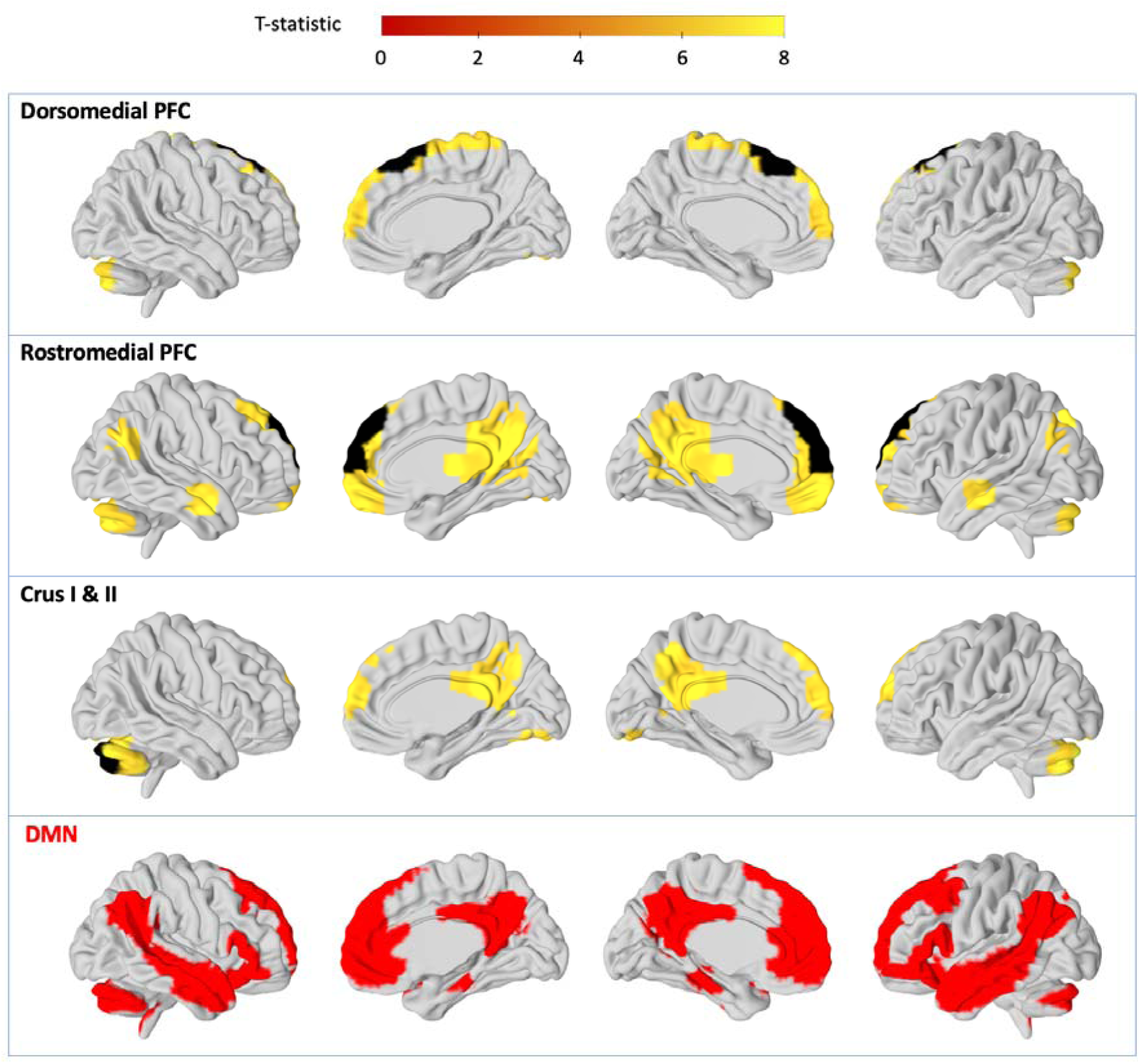
Intrinsic connectivity of the significant brain clusters involved in *Goal-directed Remoteness*. The first three rows show, for each significant cluster from the task-based fMRI analysis (displayed in black), the results of the whole-brain one-sample t-test of seed-to-voxel resting state connectivity (displayed in yellow to orange). Statistical analyses were thresholded for significance at *P_FWE_* < 0.05 at the voxel level with a spatial cluster extent threshold of 10 contiguous voxels. The fourth row shows a representation of the default mode network (DMN).(Yeo et al. 2011; Buckner et al. 2011; Choi et al. 2012) dorsomedial PFC: dorsomedial prefrontal cortex; rostromedial PFC: rostromedial prefrontal cortex

**Table 2.**
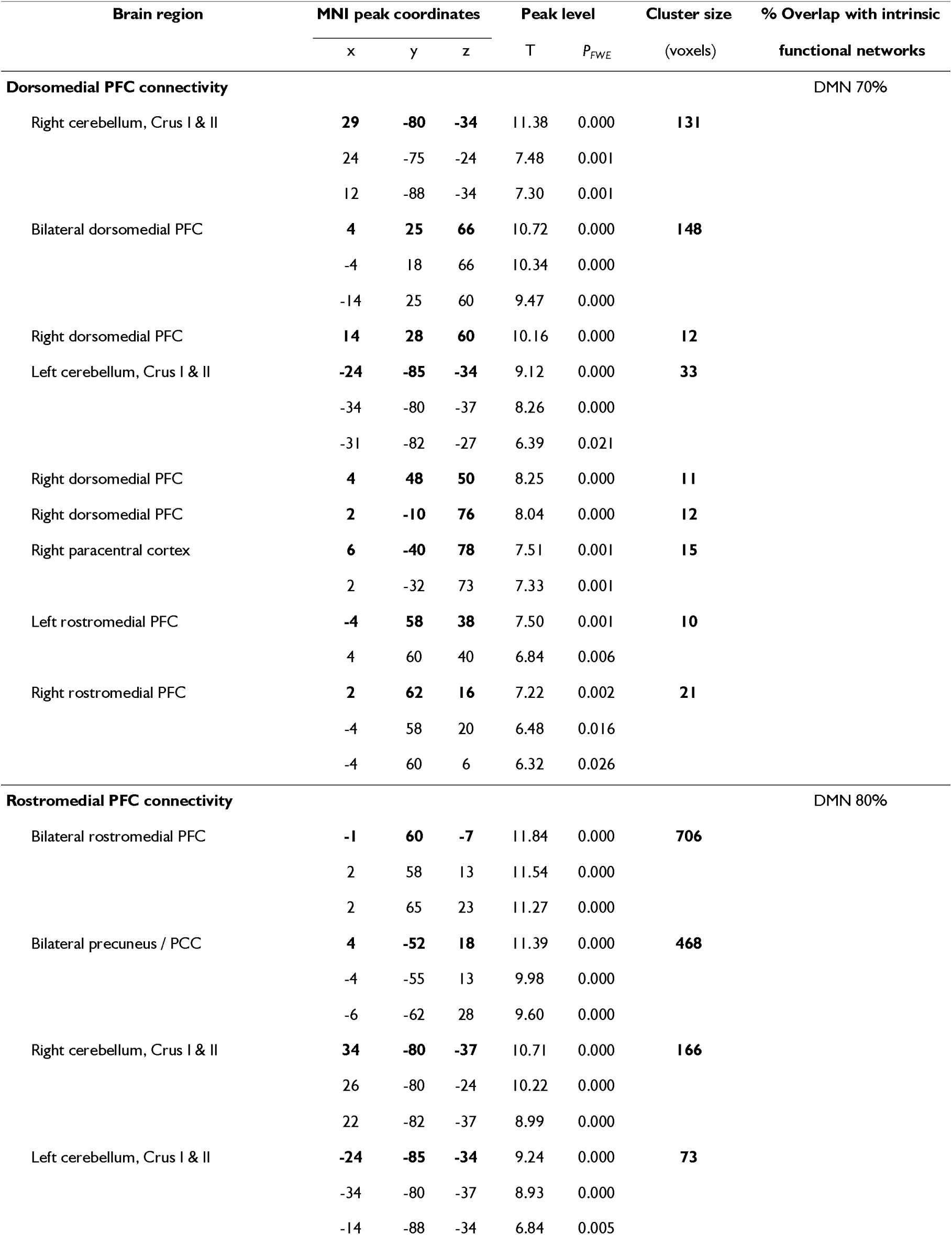
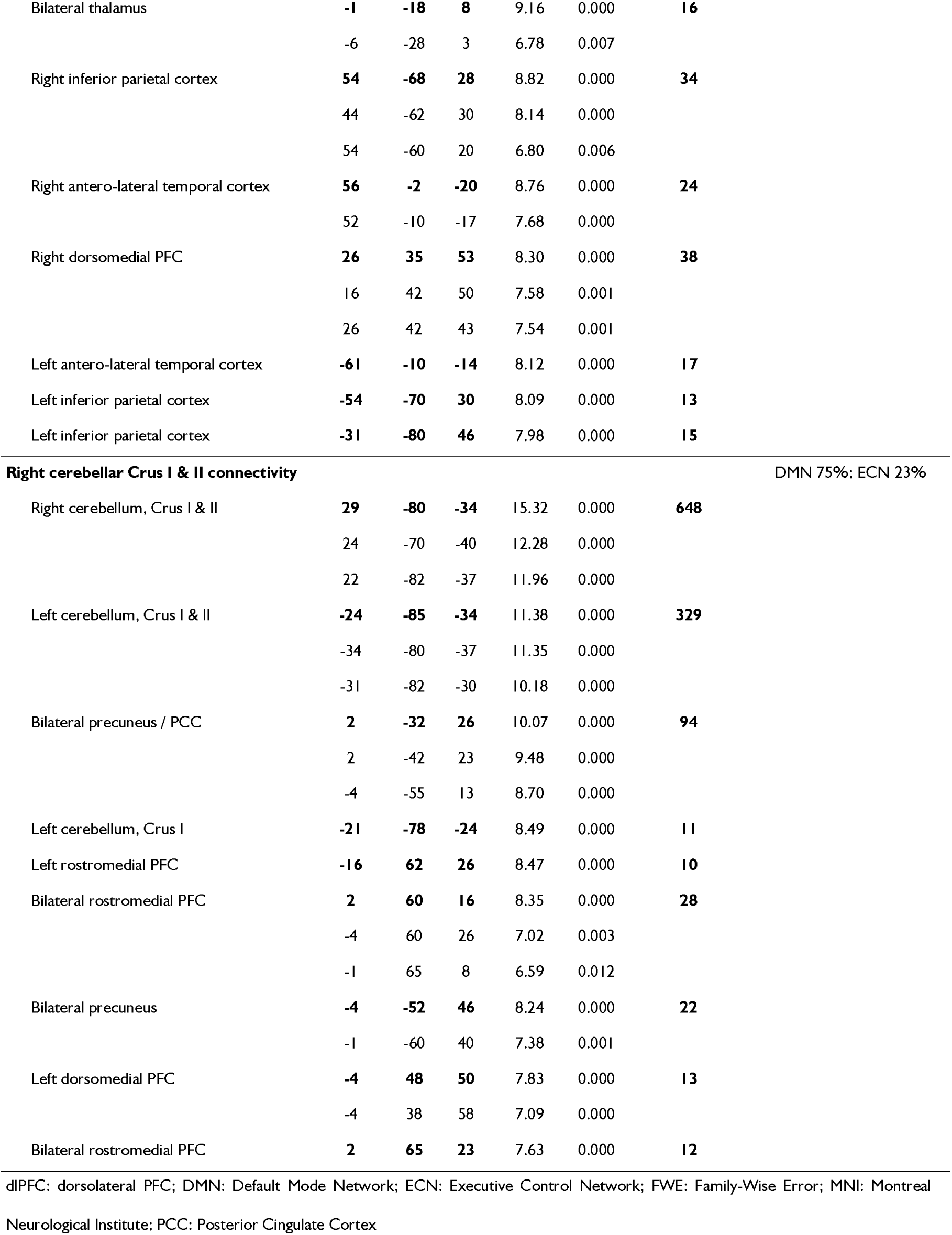
Whole-brain one-sample t-tests of seed-to-voxel resting state connectivity for each Goal-directed Remoteness-related region. Significant clusters for whole-brain one-sample t-tests of each Goal-directed Remoteness-related region’s resting state connectivity are reported, including MNI coordinates, and T-score of significant clusters’ maxima, cluster size, and the overlaps of each connectivity map with Yeo and colleagues’ intrinsic functional networks(Yeo et al. 2011; Buckner et al. 2011; Choi et al. 2012) (overlaps above 10% are indicated). Statistical analyses were thresholded for significance at PFWE-corrected < .05 at the voxel level, with a spatial cluster extent threshold of 10 contiguous voxels.

## Discussion

The current study refines the characterization of creative neurocognition by identifying key brain regions involved in goal-directed remote thinking, a fundamental aspect of creativity.(Beaty et Kenett 2023) Using a simple word-to-word association task, the FGAT, we isolate goal-directed from spontaneous remote thinking and identify a prefronto-cerebellar network involved in the goal-directed generation of remote semantic associations. It comprises interconnected rostromedial PFC, dorsomedial PFC, and cerebellar Crus I & II regions, which are embedded in and connected with the DMN. Our results question the classical view associating the DMN with spontaneous cognitive processes(Raichle et al. 2001; Anticevic et al. 2012), suggesting that the role of the DMN in creativity is more complex than initially thought and may involve functional subnetworks. Thus, our findings help clarify which role DMN regions can play in creative thinking.(Beaty et al. 2019; Bendetowicz et al. 2018; Beaty, Kenett, et al. 2018; Beaty et al. 2016; 2015; Shofty et al. 2022; Peña et al. 2024)

More specifically, we revealed key interconnected nodes within the DMN – the rostromedial PFC, dorsomedial PFC and cerebellar Crus I & II – that were activated proportionally to *Goal-directed Remoteness* during the FGAT-Distant. The rostromedial PFC is a well-established core node of the DMN(Andrews-Hanna et al. 2014) as confirmed by its connectivity with DMN regions in the present study. Its involvement in *Goal-directed Remoteness* aligns with a growing number of studies involving healthy subjects(Green et al. 2015; Gonen-Yaacovi et al. 2013) or patients(Shamay-Tsoory et al. 2011; Bendetowicz et al. 2018; de Souza et al. 2010), highlighting the role of the rostromedial PFC (or medial frontal pole) in the production of uncommon or distant semantic associations. In particular, our results replicate previous findings from our lab showing the critical role of the right rostromedial PFC for goal-directed remote thinking measured using the same FGAT task in patients with focal brain lesions (Bendetowicz et al. 2018) or neurodegenerative diseases.(Altmayer et al. 2024; Altmayer, Ovando-Tellez, Bieth, Batrancourt, Rametti-Lacroux, Moreno-Rodriguez, et al. 2025) Using a comparable verb-generation task, Green *et al*.(Green et al. 2015) also demonstrated the recruitment of the left rostromedial PFC to generate remote semantic associations. The critical role of the rostromedial PFC is further supported by studies that reported an increased remoteness of verbal associations after rostromedial PFC excitation using transcranial direct-current stimulation.(Green et al. 2017; Brunyé et al. 2015) However, in these studies, the distance of spontaneous associations was not taken into account, conflating spontaneous and goal-directed remoteness. Our results go beyond this literature by isolating the contribution of the rostromedial PFC to thinking away from common, spontaneous associations, thus clarifying its involvement in the goal-directed process of remote thinking.

The involvement of the rostromedial PFC in goal-oriented remote thinking may be interpreted in the light of previous studies highlighting its role in higher-level integrative functions. Research has linked this region to the integration of diverse types of memory information (Lanzoni et al. 2020; Murphy et al. 2018), as well as to scheme-mediated memory retrieval, i.e., the activation of superordinate knowledge structures that reflect abstracted, generalized commonalities across multiple experiences.(Gilboa et Marlatte 2017; Lagarde et al. 2015) This form of memory retrieval is believed to support processes such as analogical reasoning(Gick et Holyoak 1983) and future thinking, which involves using both past episodic memories and semantic knowledge to envision potential future scenarios(Schacter et al. 2007; Irish et al. 2012; Eustache et al. 2016), both of which are considered fundamental components of creative thought.(Green et al. 2012; Beaty, Thakral, et al. 2018; Thakral et al. 2021)

The dorsomedial PFC, also activated during goal-directed remote thinking, is a key region of the dorsomedial DMN subsystem related to semantic control(Andrews-Hanna et al. 2014), as confirmed by its connectivity profile in the current study. The dorsomedial PFC is also a central region of the semantic control network regulating the access to semantic knowledge and its manipulation based on task or context demands(Jackson 2021; Lambon Ralph et al. 2017), which has been linked with creativity.(Krieger-Redwood et al. 2023; Gonen-Yaacovi et al. 2013; Boccia et al. 2015) Thought to signal a need for behavioral adaptation(Clairis et Lopez-Persem 2023), the dorsomedial PFC may facilitate shifts in mental representations by fostering flexible and adaptive access to several semantic options, and contributing to bypass dominant, conventional semantic associations to generate creative ones.(Krieger-Redwood et al. 2023; Altmayer et al. 2024; Alario et al. 2006) Finally, the dorsomedial PFC in the DMN is recognized for its role in mentalization(Andrews-Hanna et al. 2014), i.e., in the metacognitive process of reflecting on one’s own mental states. Such processes may play an important role in remote thinking, for instance to monitor how unique (unlikely to occur to others) an evoked association may be.(Lebuda et Benedek 2025; Jia et al. 2019) However, future studies are needed to clarify the exact cognitive contribution of the dorsomedial PFC in remote thinking.

Often overlooked in creativity research, cerebellar regions encompassing the right Crus I & II were also found to be activated proportionally to *Goal-directed Remoteness*. The cerebellum, in particular Crus I & II, has been linked to language and semantics(Ito 2008; Stoodley et Schmahmann 2009; Turker et al. 2023; Kubinec et al. 2026), and to both verbal and non-verbal creativity.(Gonen-Yaacovi et al. 2013; Cogdell-Brooke et al. 2020; Gao et al. 2020) Our RSFC findings revealed that these regions were primarily connected to the DMN, but also to the ECN.(Buckner et al. 2011) This connectivity positions Crus I & II at the intersection of DMN and ECN and might explain its previously described involvement in both automatic uncontrolled processes (such as the retrieval and prediction of words’ dominant associates(Kubinec et al. 2026; Petríková et al. 2023; Lesage et al. 2017; D’Mello et al. 2017)) and higher-level semantic control processes (such as conceptual integration and representation(LeBel et D’Mello 2023; LeBel et al. 2021), semantic relatedness judgement(Gatti et al. 2020; Herault et al. 2024) and verbal working memory(Stoodley et Schmahmann 2009; Ito 2008). Our results provide evidence for the role of Crus I & II in thinking away from dominant spontaneous associates. Interestingly, we also found that the correlation between right Crus I & II activity and *Goal-directed Remoteness* diminished when controlling for cue-words’ steepness, indicating that their engagement is modulated by the need to override strong associative responses, rather than a function of semantic remoteness. Given the cerebellum’s role in predictive modelling and error detection(Ito 2008; Koziol et al. 2014; Sokolov et al. 2017; Lesage et al. 2017), the increased cerebellar activation with steeper cue-words might reflect the monitoring of deviations from internal predictions, i.e., from contextually dominant, most adequate, semantic associations.(Petríková et al. 2023; Lesage et al. 2017; D’Mello et al. 2017) In creative contexts, this may involve monitoring cue-words’ dominant semantic associates from which to think away, to allow individuals to think remotely, creatively.(Cogdell-Brooke et al. 2020; Sunavsky et Poppenk 2020; Neumann et al. 2018; Gao et al. 2020) Importantly, the Crus I & II cluster identified here differs from that previously found critical for spontaneous FGAT-First responses(Altmayer et al. 2024), which was prominently connected to the ECN and overlapped with a cluster encoding the adequacy of FGAT-Distant responses in another study which used the same dataset.(Moreno-Rodriguez et al. 2025) By contrast, the current cluster is predominantly connected to the DMN and overlaps with a region encoding the originality of FGAT-Distant responses.(Moreno-Rodriguez et al. 2025) This suggests that distinct subregions within Crus I & II may support different cognitive demands during remote thinking, depending on task context and goals.(Stoodley et Schmahmann 2009; Sokolov et al. 2017; Guell et al. 2018; King et al. 2019)

Overall, our findings challenge the assumption that the DMN selectively supports passive, spontaneous aspects of creative cognition, pointing out the need to reframe the role of the DMN (or of its subnetworks) in creativity. Indeed, our results reveal the central role of a prefronto-cerebellar subnetwork overlapping with the DMN in goal-directed forms of remote thinking (with the goal of thinking remotely, creatively). While goal-directed cognition is classically associated with the ECN(Benedek et Jauk 2018), our study aligns with studies suggesting that DMN and ECN are jointly engaged during creative thinking.(Beaty, Kenett, et al. 2018; Beaty et al. 2015; 2016) The classical description of the DMN as the support of spontaneous, uncontrolled cognition, largely comes from its initial description in task-free experimental paradigms, and its frequent deactivation during externally focused tasks.(Raichle et al. 2001; Anticevic et al. 2012) However, our results support the alternative view that part of the DMN, rather than functioning as a strictly “task-negative” network, can also play a role in active processes involved in goal-directed remote thinking.

Such finding aligns with an emerging view of the DMN as supporting the retrieval of both semantic and episodic memory information(Andrews-Hanna et al. 2014; Kim 2016; Irish et Vatansever 2020; Beaty et al. 2020), their integration(Fernandino et Binder 2024; Lanzoni et al. 2020), and the self-generation of mental contents.(Zabelina et Andrews-Hanna 2016; Smallwood et al. 2021; Murphy et al. 2018) The DMN may integrate information from various modalities of memory traces, including semantic, episodic, emotional, or reward information, to generate internal representations that help us simulate and explore complex scenarios even in the absence of external input.(Fernandino et Binder 2024) This integrative function of the DMN is likely central to creative thinking, enabling the connection of remote elements of knowledge in novel ways, as in remote thinking. Furthermore, recent evidence suggests that the DMN may contribute to goal-directed aspects of semantic exploration(Bendetowicz et al. 2018; Wang et al. 2021; Beaty et al. 2015; Beaty, Kenett, et al. 2018), during which it may contribute to representing the semantic goal.(Wang et al. 2021) Finally, while the DMN is typically associated with creative idea generation, it has recently been shown to also play a role in evaluating the relatedness of semantic associations(Herault et al. 2024) and monitoring their originality during goal-directed remote thinking(Moreno-Rodriguez et al. 2025), thereby ensuring adaptation to the creative goal. Hence, the DMN regions we observe might not only reflect generative processes, but also the monitoring of goal-relevant information, i.e., originality.(Lopez-Persem, Mandonnet, et al. 2024; Bartoli et al. 2024)

## Limitations and future directions

While our study provides important insights into the role of the DMN in goal-directed creative thinking, several limitations should be considered. First, the use of a single verbal task to assess goal-directed creative cognition may limit the generalizability of our findings to other forms of creative cognition, such as non-verbal creativity. Also, task-fMRI analyses are correlational methods that limit causal conclusions. However, our results replicate findings obtained with the same task in brain-lesioned patients with frontotemporal dementia, which support the critical role of the rostromedial PFC and dorsomedial PFC for the intentional generation of remote semantic associates.(Altmayer et al. 2024; Altmayer, Ovando-Tellez, Bieth, Batrancourt, Rametti-Lacroux, Moreno-Rodriguez, et al. 2025)

Finally, it may seem surprising that we did not find a dominance of the ECN, traditionally related to goal-directed control processes, in goal-directed remote thinking. We nevertheless found the involvement of the dorsomedial PFC, a region at the intersection of the DMN and ECN associated with semantic control(Jackson 2021; Lambon Ralph et al. 2017) and in the generation of original links between semantic elements.(Krieger-Redwood et al. 2023) We also didn’t observe any involvement of the left inferior frontal gyrus, another region sitting at the intersection of the DMN and ECN and classically reported in tasks involving semantic control.(Jackson 2021; Lambon Ralph et al. 2017) It is possible that some brain regions involved in executive or semantic control processes are required to perform the FGAT-Distant condition but were not observed using parametric modulation because their recruitment is not proportional to the achieved distance in the response. Hence, our results don’t exclude the role of executive and semantic control processes in goal-directed remote thinking; but rather highlight that, even in a goal-directed task, DMN activation correlates with the remoteness of the generated semantic associations.

Overall, our findings reveal that the DMN plays an active role in the intentional generation of remote associations between distant concepts. They identify a prefronto-cerebellar DMN subnetwork supporting goal-directed remote thinking, including the rostromedial PFC, the dorsomedial PFC, and Crus I & II of the cerebellum. These results open an avenue for future research, intending to decipher the exact computations performed by these DMN regions to enable goal-directed remote thinking.

## Acknowledgements

We thankfully acknowledge the contributions the PRISME and CENIR ICM platforms for their help in collecting cognitive and brain imaging data, respectively.

Finally, we sincerely thank the volunteers for participating in this study.

## Supplementary material

Supplementary material is available online.

