## Supplementary Material for "From default to creativity: prefrontal and cerebellar contributions of the default mode network to goal-directed remote thinking"

**Authors affiliations**:

^1^ FrontLab, Sorbonne University, Institut du Cerveau – Paris Brain Institute – ICM, Inserm, CNRS, AP-HP, Hôpital de la Pitié Salpêtrière, Paris, France

^2^ AP-HP, Groupe Hospitalier Pitié-Salpêtrière, Department of Neurology, IM2A, Paris, France

^3^ CENIR, Institut du Cerveau – Paris Brain Institute – ICM, Inserm, CNRS, AP-HP, Hôpital de la Pitié Salpêtrière, Paris, France.

### **Supplementary Tables**

**Supplementary Table 1.**

**Cue-words’ lexical and semantic characteristics**

For each cue-word are provided: movies-based, books-based and average lexical frequencies, computed with Lexique 3.8(New et al. 2004); [www.lexique.org](http://www.lexique.org); semantic richness computed as the ratio between the number of unique semantic associates for this cue-word and the total number of responses given for this cue-word, based on the French free-association norms(Debrenne 2011); and steepness computed as the frequency ratio between the most frequent and the second most frequent associates of a given cue-word, based on the French free-association norms.

| **Cueword**  (in French) | **Cueword**  (in English) | **Lexical frequency**  **Movies** | **Lexical frequency**  **Books** | **Lexical frequency**  **Average** | **Semantic richness** | **Steepness** |
| --- | --- | --- | --- | --- | --- | --- |
| acte | action | 39,19 | 35,88 | 37,54 | 23,93 | 2,08 |
| appel | call | 80,88 | 56,69 | 68,79 | 18,90 | 5,98 |
| arbre | tree | 49,29 | 67,16 | 58,23 | 17,20 | 1,72 |
| avis | opinion | 139,22 | 65,14 | 102,18 | 22,99 | 4,52 |
| cause | cause | 213,51 | 188,04 | 200,78 | 16,97 | 1,14 |
| chambre | bedroom | 263,93 | 380,07 | 322,00 | 17,92 | 4,04 |
| chien | dog | 158,77 | 117,64 | 138,21 | 12,83 | 14,73 |
| colline | hill | 15,61 | 30,07 | 22,84 | 16,39 | 6,06 |
| début | start | 109,88 | 128,51 | 119,20 | 9,18 | 8,44 |
| désir | desire | 31,59 | 96,69 | 64,14 | 18,79 | 1,72 |
| direction | direction | 43,25 | 103,04 | 73,15 | 22,06 | 5,74 |
| doigt | finger | 39,83 | 80,34 | 60,09 | 13,66 | 11,15 |
| école | school | 197,04 | 128,51 | 162,78 | 24,22 | 1,11 |
| enfant | child | 287,26 | 381,96 | 334,61 | 27,94 | 1,10 |
| façon | manner | 212,60 | 259,26 | 235,93 | 11,38 | 24,08 |
| faim | hunger | 127,49 | 74,93 | 101,21 | 18,28 | 1,03 |
| fil | thread | 51,83 | 75,95 | 63,89 | 19,51 | 2,54 |
| flamme | flame | 12,61 | 38,38 | 25,50 | 10,33 | 20,59 |
| garçon | boy | 188,41 | 186,96 | 187,69 | 13,25 | 17,48 |
| gauche | left | 71,66 | 133,78 | 102,72 | 9,07 | 13,81 |
| genre | gender | 219,66 | 155,20 | 187,43 | 15,55 | 1,05 |
| habitude | habit | 89,71 | 128,51 | 109,11 | 25,56 | 1,23 |
| herbe | grass | 27,64 | 86,08 | 56,86 | 19,22 | 1,36 |
| horizon | horizon | 7,80 | 61,08 | 34,44 | 15,48 | 1,59 |
| jambe | leg | 46,31 | 49,93 | 48,12 | 19,13 | 1,35 |
| jardin | garden | 54,01 | 148,72 | 101,37 | 22,84 | 1,29 |
| jeunesse | youth | 27,21 | 83,24 | 55,23 | 29,51 | 5,95 |
| larme | tear | 5,15 | 10,81 | 7,98 | 10,62 | 1,19 |
| lèvre | lip | 4,00 | 20,74 | 12,37 | 14,71 | 5,88 |
| lit | bed | 176,10 | 315,74 | 245,92 | 15,80 | 7,29 |
| lune | moon | 58,29 | 63,24 | 60,77 | 12,82 | 2,47 |
| lutte | struggle | 22,00 | 37,36 | 29,68 | 22,49 | 4,94 |
| machine | machine | 44,94 | 58,99 | 51,97 | 25,62 | 1,76 |
| mari | husband | 254,92 | 118,38 | 186,65 | 10,46 | 7,51 |
| masse | mass | 11,64 | 60,54 | 36,09 | 25,96 | 9,20 |
| mer | sea | 99,49 | 246,55 | 173,02 | 16,80 | 1,96 |
| morceau | piece | 40,65 | 57,84 | 49,25 | 13,97 | 1,67 |
| mur | wall | 58,90 | 172,57 | 115,74 | 25,74 | 1,73 |
| nord | north | 50,38 | 72,30 | 61,34 | 10,96 | 8,78 |
| objet | object | 26,88 | 67,09 | 46,99 | 29,70 | 6,82 |
| oncle | uncle | 124,11 | 121,96 | 123,04 | 8,48 | 5,75 |
| page | page | 25,16 | 55,88 | 40,52 | 13,68 | 4,74 |
| passage | passage | 40,20 | 136,82 | 88,51 | 23,98 | 2,55 |
| passion | passion | 30,71 | 68,72 | 49,72 | 22,74 | 4,09 |
| peau | skin | 83,83 | 174,26 | 129,05 | 29,50 | 1,58 |
| pierre | stone | 40,58 | 119,39 | 79,99 | 20,86 | 2,09 |
| position | position | 55,24 | 53,85 | 54,55 | 33,06 | 1,34 |
| preuve | proof | 60,79 | 50,54 | 55,67 | 29,78 | 1,18 |
| principe | principle | 15,62 | 30,88 | 23,25 | 34,32 | 1,30 |
| question | question | 293,63 | 232,50 | 263,07 | 9,68 | 7,90 |
| rapport | report | 115,57 | 87,23 | 101,40 | 27,20 | 1,56 |
| rayon | ray | 19,32 | 27,03 | 23,18 | 15,08 | 4,32 |
| rôle | role | 61,20 | 88,51 | 74,86 | 13,71 | 1,38 |
| rose | rose | 18,43 | 66,62 | 42,53 | 24,13 | 5,31 |
| science | science | 25,28 | 24,93 | 25,11 | 30,47 | 1,14 |
| sens | sense | 117,57 | 217,50 | 167,54 | 25,81 | 4,07 |
| soeur | sister | 155,22 | 116,55 | 135,89 | 15,22 | 6,90 |
| table | table | 111,44 | 341,08 | 226,26 | 14,42 | 4,00 |
| toit | roof | 42,63 | 54,59 | 48,61 | 13,81 | 6,95 |
| vache | cow | 36,24 | 26,08 | 31,16 | 14,48 | 4,04 |
| vin | wine | 80,92 | 99,93 | 90,43 | 20,00 | 1,55 |
| visage | face | 125,52 | 490,54 | 308,03 | 22,93 | 1,45 |

**Supplementary Table 2.**

**Results for linear mixed-effects models**

Separate linear mixed-effects models with random effects on both individuals and cue words were used to explore whether cue-words’ semantic richness, steepness, or trial response time predicted *Goal-directed Remoteness*.

| ***Goal-Directed Remoteness* ~ Semantic richness + (Semantic richness \| Cue-Word) + (Semantic richness \| Participant)** | | | | |
| --- | --- | --- | --- | --- |
| Fixed Effects |  |  |  |  |
|  | Estimate | Standard Error | Degrees of freedom | *t-*value (*P*) |
| (Intercept) | 4.22^-1^ | 5.50^-2^ | 53.41 | 7.48 (*P* < .001) *** |
| Semantic richness | -1.36^-2^ | 2.46^-3^ | 28.00 | -5.54 (*P* < .001) *** |
| Random Effects |  |  |  |  |
| Groups | Name | Variance | Standard deviation | Correlation |
| Cue-word | (Intercept) | 6.56^-2^ | 2.56^-1^ |  |
|  | Semantic richness | 1.22^-4^ | 1.10^-2^ | -0.96 |
| Participant | (Intercept) | 1.88^-2^ | 1.37^-1^ |  |
|  | Semantic richness | 1.69^-5^ | 4.11^-3^ | -0.98 |
| Residual |  | 5.20^-2^ | 2.28^-1^ |  |
| ***Goal-Directed Remoteness* ~ Steepness + (Steepness \| Cue-Word) + (Steepness \| Participant)** | | | | |
| Fixed Effects |  |  |  |  |
|  | Estimate | Standard Error | Degrees of freedom | *t-*value (*P*) |
| (Intercept) | 7.00^-2^ | 2.36^-2^ | 52.68 | 2.97 (*P* = .004) ** |
| Steepness | 1.99^-2^ | 6.24^-3^ | 24.68 | 3.19 (*P* = .004) ** |
| Random Effects |  |  |  |  |
| Groups | Name | Variance | Standard deviation | Correlation |
| Cue-word | (Intercept) | 1.16^-2^ | 1.08^-1^ |  |
|  | Steepness | 8.60^-4^ | 2.93^-2^ | -0.72 |
| Participant | (Intercept) | 2.69^-3^ | 5.19^-2^ |  |
|  | Steepness | 3.13^-5^ | 5.59^-3^ | 0.05 |
| Residual |  | 5.19^-2^ | 2.28^-1^ |  |
| ***Goal-Directed Remoteness* ~ Response Time + (Response Time \| Cue-Word) + (Response Time \| Participant)** | | | | |
| Fixed Effects |  |  |  |  |
|  | Estimate | Standard Error | Degrees of freedom | *t-*value (*P*) |
| (Intercept) | 1.72^-1^ | 2.77^-2^ | 69.34 | 6.19 (*P* < .001) *** |
| Response Time | 8.88^-7^ | 2.05^-6^ | 37.94 | 0.43 (*P* = .668) |
| Random Effects |  |  |  |  |
| Groups | Name | Variance | Standard deviation | Correlation |
| Cue-word | (Intercept) | 2.55^-2^ | 1.58^-1^ |  |
|  | Response Time | 9.00^-14^ | 3.00^-7^ | 1.00 |
| Participant | (Intercept) | 8.79^-3^ | 9.38^-2^ |  |
|  | Response Time | 6.97^-11^ | 8.35^-6^ | -0.83 |
| Residual |  | 5.19^-2^ | 2.28^-1^ |  |

**Supplementary Table 3.**

**Significant peak coordinates of the whole-brain parametric modulation by *Goal-directed Remoteness* orthogonalized on cue-words’ semantic richness**

Significant clusters from the whole-brain fMRI analysis of the FGAT-Distant task with *Goal-directed Remoteness* orthogonalized on cue-words’ semantic richness as parametric modulator are reported, including MNI coordinates, T-score and *P_uncorrected_* value of significant clusters’ maxima, cluster size, cluster *P_FWE_* value, and related Brodmann areas. Statistical analyses were thresholded for significance at *P_uncorrected_* < 0.001 at the voxel level and *P_FWE_* < 0.05 at the cluster level.

| **Brain region** | **BA** | **MNI peak coordinates** | | | **Peak level** | | **Cluster level** | |
| --- | --- | --- | --- | --- | --- | --- | --- | --- |
|  |  | x | y | z | T | *P_uncorrected_* | voxels | *P_FWE_* |
| Bilateral dorsomedial and | 10, 9, 6, 8 | **-11** | **25** | **58** | 5.75 | 0.000 | **691** | **0.000** |
| rostromedial PFC |  | -8 | 62 | 23 | 5.12 | 0.000 |  |  |
|  |  | 6 | 22 | 63 | 4.87 | 0.000 |  |  |
| Right cerebellum, Crus I & II | NA | **26** | **-80** | **-37** | 5.28 | 0.000 | **158** | **0.016** |

BA: Brodmann Areas; FWE: Family Wise Error; RSFC: Resting-State Functional Connectivity; MNI: Montreal Neurological Institute; NA: Not Applicable; PFC: Prefrontal Cortex

**Supplementary Table 4.**

**Significant peak coordinates of the whole-brain parametric modulation by *Goal-directed Remoteness* orthogonalized on cue-words’ steepness**

Significant clusters from the whole-brain fMRI analysis of the FGAT-Distant task with *Goal-directed Remoteness* orthogonalized on cue-words’ steepness as parametric modulator are reported, including MNI coordinates, T-score and *P_uncorrected_* value of significant clusters’ maxima, cluster size, cluster *P_FWE_* value, and related Brodmann areas. Statistical analyses were thresholded for significance at *P_uncorrected_* < 0.001 at the voxel level and *P_FWE_* < 0.05 at the cluster level.

| **Brain region** | **BA** | **MNI peak coordinates** | | | **Peak level** | | **Cluster level** | |
| --- | --- | --- | --- | --- | --- | --- | --- | --- |
|  |  | x | y | z | T | *P_uncorrected_* | voxels | *P_FWE_* |
| Bilateral dorsomedial PFC | 6 | **-8** | **25** | **60** | 5.47 | 0.000 | **225** | **0.003** |
|  |  | 12 | 30 | 58 | 4.65 | 0.000 |  |  |
|  |  | 6 | 25 | 63 | 4.48 | 0.000 |  |  |
| Bilateral rostromedial PFC | 10, 9 | **-8** | **65** | **23** | 5.44 | 0.000 | **275** | **0.001** |
|  |  | -6 | 55 | 38 | 4.52 | 0.000 |  |  |
|  |  | -4 | 48 | 46 | 4.15 | 0.000 |  |  |

BA: Brodmann Areas; FWE: Family Wise Error; RSFC: Resting-State Functional Connectivity; MNI: Montreal Neurological Institute; NA: Not Applicable; PFC: Prefrontal Cortex

**Supplementary Table 5.**

**Significant peak coordinates of the whole-brain parametric modulation by *Goal-directed Remoteness* orthogonalized on response time**

Significant clusters from the whole-brain fMRI analysis of the FGAT-Distant task with *Goal-directed Remoteness* orthogonalized on FGAT-Distant response time as parametric modulator are reported, including MNI coordinates, T-score and *P_uncorrected_* value of significant clusters’ maxima, cluster size, cluster *P_FWE_* value, and related Brodmann areas. Statistical analyses were thresholded for significance at *P_uncorrected_* < 0.001 at the voxel level and *P_FWE_* < 0.05 at the cluster level.

| **Brain region** | **BA** | **MNI peak coordinates** | | | **Peak level** | | **Cluster level** | |
| --- | --- | --- | --- | --- | --- | --- | --- | --- |
|  |  | x | y | z | T | *P_uncorrected_* | voxels | *P_FWE_* |
| Bilateral dorsomedial and | 6, 10, 9, 8 | **-11** | **25** | **58** | 5.82 | 0.000 | **554** | **0.000** |
| rostromedial PFC |  | -8 | 62 | 26 | 5.00 | 0.000 |  |  |
|  |  | 6 | 22 | 63 | 4.90 | 0.000 |  |  |
| Right cerebellum, Crus I & II | NA | **29** | **-78** | **-37** | 5.34 | 0.000 | **115** | **0.049** |

BA: Brodmann Areas; FWE: Family Wise Error; RSFC: Resting-State Functional Connectivity; MNI: Montreal Neurological Institute; NA: Not Applicable; PFC: Prefrontal Cortex

### **Supplementary Figures**

**Supplementary** **Figure 1. Distribution of the semantic similarity with the cue-word in FGAT-First and FGAT-Distant conditions.**

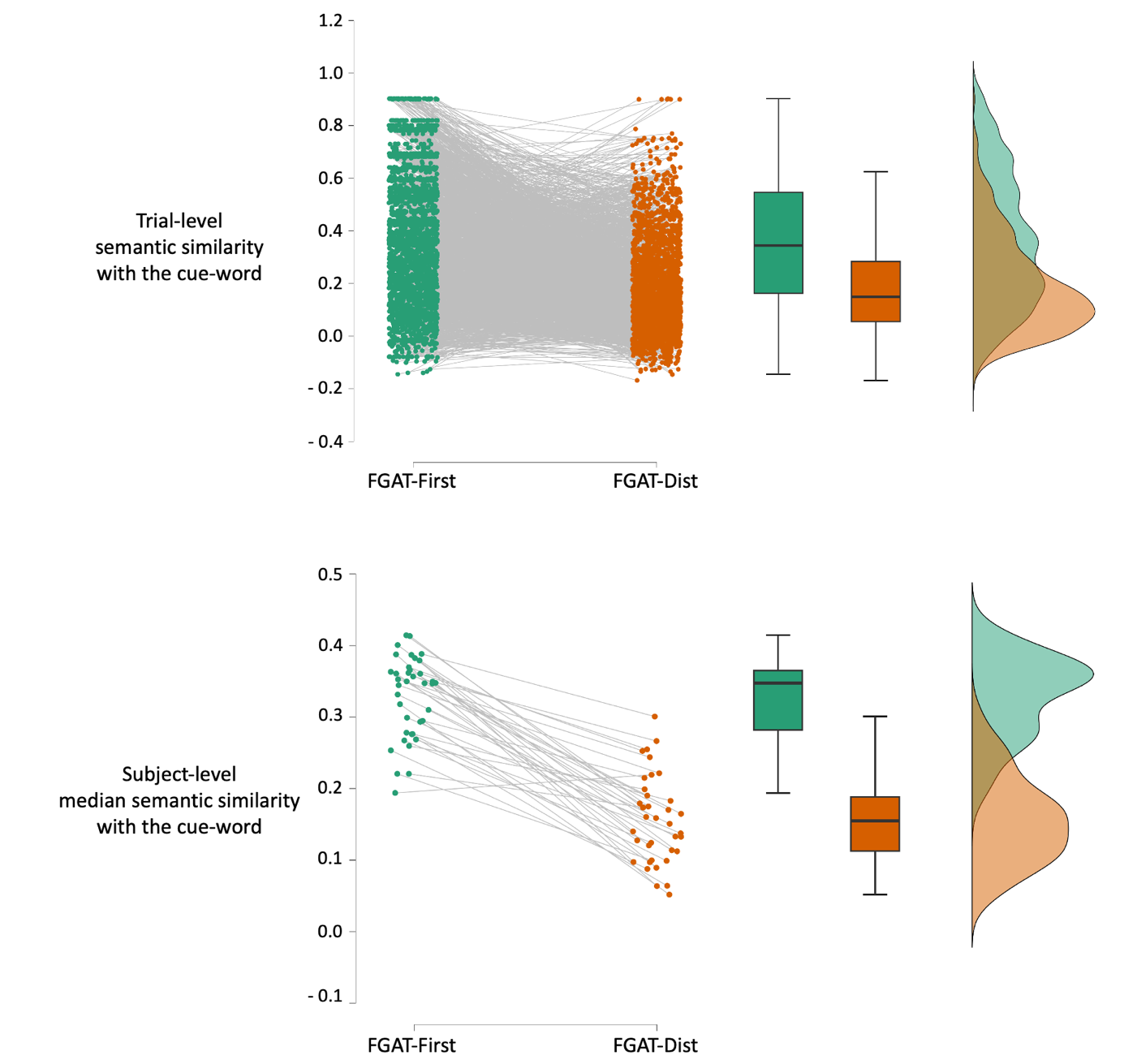

Distributions of semantic similarity between cue words and responses in the FGAT-First and FGAT-Distant conditions are displayed. The top panel shows trial-level semantic similarity values for all responses, used in the pairwise comparison between the two conditions, illustrating significantly higher semantic similarities in the FGAT-First than in the FGAT-Distant condition. The bottom panel displays subject-level median semantic similarities.

**Supplementary** **Figure 2. FGAT-Distant brain activation related to *Goal-directed Remoteness* orthogonalized on cue-words’ semantic richness**

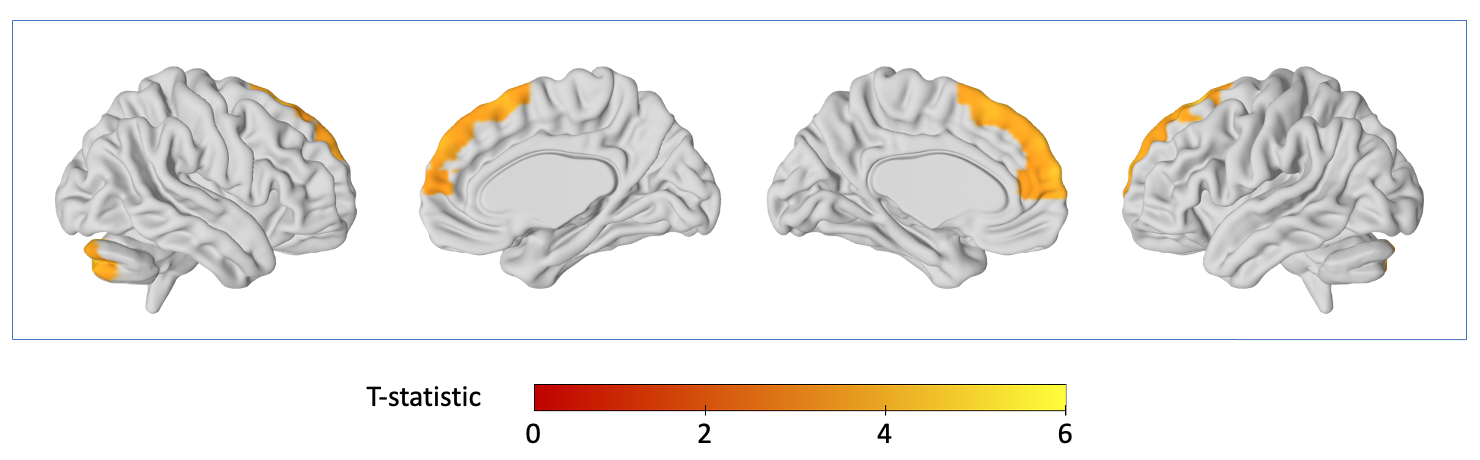

Whole-brain parametric modulation revealing significant regions modulated by *Goal-directed Remoteness* orthogonalized on cue-words’ semantic richness during FGAT distant. Significant clusters are shown in colour. Statistical analyses were thresholded for significance at *P_FWE_* < 0.05 at the cluster level with a spatial cluster extent threshold of 10 contiguous voxels.

**Supplementary** **Figure 3. FGAT-Distant brain activation related to *Goal-directed Remoteness* orthogonalized on cue-words’ steepness**

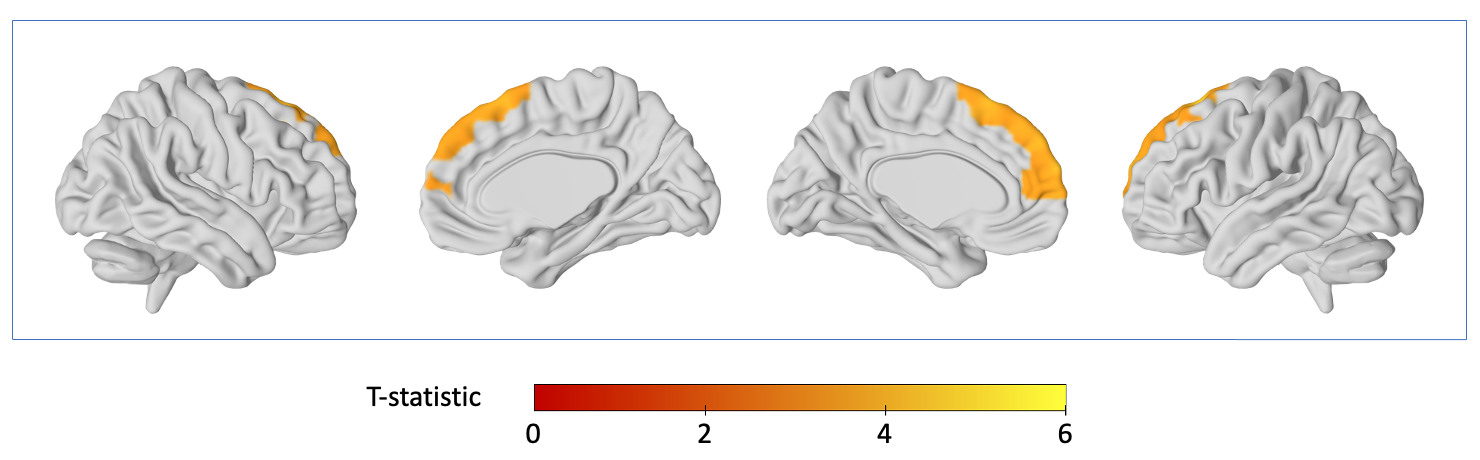

Whole-brain parametric modulation revealing significant regions modulated by *Goal-directed Remoteness* orthogonalized on cue-words’ steepness during FGAT distant. Significant clusters are shown in colour. Statistical analyses were thresholded for significance at *P_FWE_* < 0.05 at the cluster level with a spatial cluster extent threshold of 10 contiguous voxels.

**Supplementary** **Figure 4. FGAT-Distant brain activation related to *Goal-directed Remoteness* orthogonalized on trials’ response time**

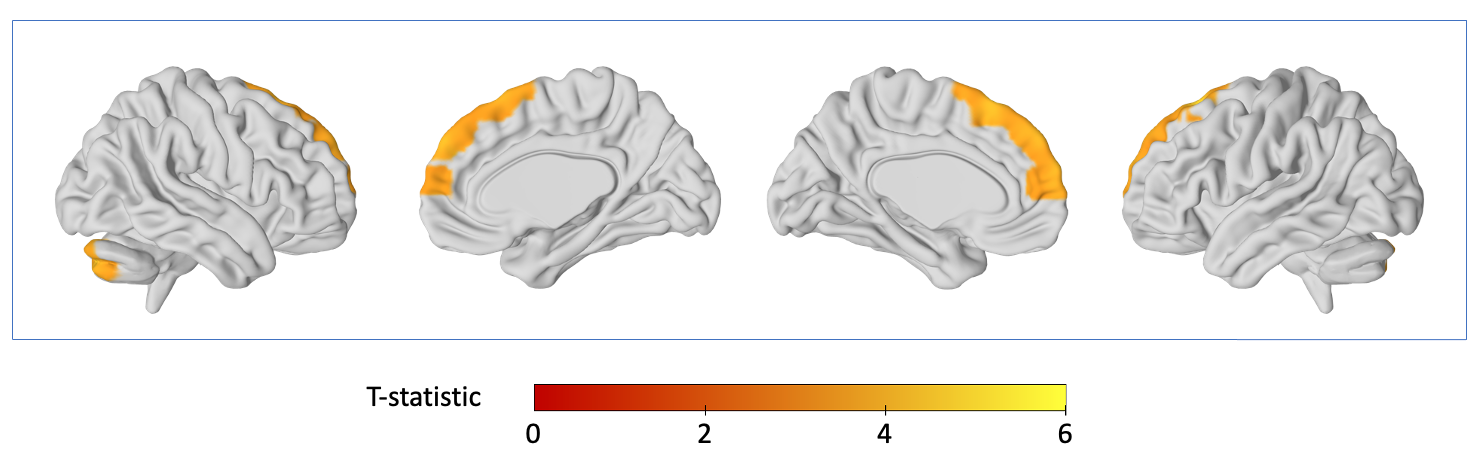

Whole-brain parametric modulation revealing significant regions modulated by *Goal-directed Remoteness* orthogonalized on trials’ response time during FGAT distant. Significant clusters are shown in colour. Statistical analyses were thresholded for significance at *P_FWE_* < 0.05 at the cluster level with a spatial cluster extent threshold of 10 contiguous voxels.

### **Supplementary References**

Debrenne, Michèle. 2011. *Le dictionnaire des associations verbales du français et ses applications*.

New, Boris, Christophe Pallier, Marc Brysbaert, et Ludovic Ferrand. 2004. « Lexique 2 : A New French Lexical Database ». *Behavior Research Methods, Instruments, & Computers* 36 (3): 516‑24. https://doi.org/10.3758/BF03195598.
